# Sex-specific lipid profiles in the muscle of Atlantic salmon juveniles

**DOI:** 10.1101/2020.11.13.379321

**Authors:** Andrew H. House, Paul V. Debes, Johanna Kurko, Jaakko Erkinaro, Reijo Käkelä, Craig R. Primmer

**Author notes:** **Present Address:** Viikinkaari 9 (PL 56), 00790 Helsinki, Finland. **Highlights:** - Atlantic salmon males showed higher concentrations of several lipid classes in muscle tissue compared to females - Muscle lipid concentrations covaried negatively with body length - Positive correlations were observed between many classes of structural and storage lipids in the muscle.

## Abstract

Energy allocation in juvenile fish can have important implications for future life-history progression. Inherited and environmental factors determine when and where individuals allocate energy, and timely and sufficient energy reserves are crucial for reaching key life stages involved in the timing of maturation and sea migration. In Atlantic salmon, lipid reserves are predominantly found in the viscera and myosepta in the muscle and have been shown to play a key role in determining the timing of maturity. This life-history trait is tightly linked to fitness in many species and can be different between males and females, however, the details of relative energy allocation in juveniles of different sexes is not well understood. Therefore, the aim of this study was to investigate the effects of sex, genetics and environment during juvenile development of salmon on the amount and composition of their lipid reserves. To do so, juvenile salmon were fed one of two different lipid food contents during their first summer and autumn under common-garden conditions. Muscle lipid composition and concentrations were determined by thin layer chromatography. The muscle lipid class concentrations covaried negatively with body length and males showed higher concentrations than females for phosphatidylcholine, cholesterol, sphingomyelin, and triacylglycerol. This sex-specific difference in major lipid classes presents a new scope for understanding the regulation of lipids during juvenile development and gives direction for understanding how lipids may interact and influence major life-history traits in Atlantic salmon.

## Introduction

Optimal allocation of energy between activity, growth and storage are crucial for reaching key life stages, with survival to first reproduction being one of the primary determinants of success (Post & Parkinson, 2001). A tradeoff exists between somatic growth and energy storage, with food availability and quality contributing to this tradeoff (Mogensen & Post, 2012; Post & Parkinson, 2001). For example, fluctuating environmental conditions affect the quantity and quality of the food available for individuals living in seasonal environments, and this variability in the dietary supply of different nutrients influences how energy is allocated within their body (Post & Parkinson, 2001). Energy is mainly stored in the form of triacylglycerol (TAG), which has a higher density of energy than other biomolecules. In addition to forming the actual energy reserve, lipids have a variety of important biological roles. When released to circulation, lipids and their derivatives act as signaling molecules informing body organs and tissues on the energy status of the individual (Dupont et al., 2013; Koyama et al., 2020; Mangel and Satterthwaite, 2008; Parker and Cheung, 2020; Shalitin and Phillip, 2003; Thorpe et al., 1998). Further, the structurally diverse membrane phospholipids support the functions of integral proteins and serve as precursors for different lipid mediators modulating numerous signaling pathways (Moessinger et al., 2014; Næsje et al., 2006; Sheridan, 1988; Zhou, Ackman, & Morrison, 1996). By studying lipid class profiles, it is therefore possible to get a better understanding of the physiological status of the individual and potentially, its energetic capacity to adapt to environmental changes when passing through life stages (Murzina et al., 2016).

In Atlantic salmon, energy storage and allocation play a number of important roles controlling life-history-stage progression (Mangel and Satterthwaite, 2008; Thorpe et al., 1998). For example, there appears to be a physiological link between lipid accumulation in the body during spring and the initiation of maturation in male Atlantic salmon parr (Rowe et al., 1991). In adult Atlantic salmon, both males and females use lipid reserves for gamete production and also for reproductive activity: females for redd (nest) digging, and males for mate guarding and/or territory defense (Jonsson et al., 1997). As the production of large lipid-rich eggs is energetically more costly for females than milt production for male salmon, a sex difference exists in the energy allocated for reproduction. More specifically, females may invest 20-25% of their pre-breeding weight into gonads, whereas males may invest only 3-9% (Fleming, 1996; Jonsson et al., 1991). Further, the timing of marine migration is another life-history trait potentially influenced by lipid reserve levels (Metcalfe, 1998; Morgan et al., 2002).

Even though it has been recognized that energy allocation during salmon juvenile phases can have important implications for future sex-specific reproductive strategies (Rowe et al., 1991) or other life history decisions (Metcalfe, 1998), sex–specific juvenile energy allocation has rarely been studied. However, there has been renewed interest in studying this topic recently, due to the finding that Atlantic salmon maturation age appears to be linked with a large-effect locus, *vgll3,* that exhibits sex-dependent dominance (Barson et al., 2015). The closest associated gene *vgll3* (vestigial-like family member 3), has been shown to be a negative regulator of adipocyte production in mice (Halperin et al., 2013), and thus provides one potential genetic mechanism adjusting lipid reserve levels and consequently regulating lipid-dependent maturation in fishes, possibly in a sex-specific manner (Barson et al., 2015).

Here, we investigate sex–specific juvenile energy allocation patterns in Atlantic salmon by rearing juveniles in a common-garden setting whilst feeding two different lipid/protein ration feeds for five months and assessing lipid allocation to skeletal muscle in males and females. In addition to testing for sex differences, we further tested whether *vgll3* genotypes were linked with differences in lipid phenotypes. Our data support a sex-specific lipid strategy in juvenile Atlantic salmon and support the potential importance of levels in influencing the timing of sea migration and maturation.

## Methods

### Sampling

The Atlantic salmon juveniles used in this study derived from a first-generation hatchery stock of Atlantic salmon maintained by the Natural Resources Institute Finland (62°24ʹ50″N, 025°57ʹ15″E, Laukaa, Finland), the parents of which were caught from the Kymijoki river in southeast Finland and had been crossed in partial factorial manner. Fertilization took place in November 2016 and eggs were incubated in a flow-through incubation system at the hatchery until hatching in May 2017. Two weeks after hatching, around 2000 individuals were transferred to the Lammi Biological Station (61°04′45″N, 025°00′40″E, Lammi, Finland) for this common-garden experiment. These juveniles were reared in equal numbers in 10 flow-through tanks (63cm × 63cm × 30cm) under the local natural light cycle and temperature fluctuations of the water sourced from the nearby Lake Pääjärvi (temperature range: 6.3-17.7 °C). Fish were fed *ad libitum* with a diet consisting of fine-ground commercial fish food that differed in lipid content (high-lipid:19.9 % and low-lipid 12.6 %; Raisio Baltic Blend; Raisio Oy) for 5 months (five randomly assigned tanks per diet). In August 2017, all tank densities were reduced to 150 individuals by culling excess individuals randomly. An unplanned infection of an external parasite, *Ichthyophthirius multifiliis,* during August/September required the use of a regular formalin treatment for approximately 3 weeks to control the infection. However, high mortality was nevertheless experienced in five tanks, and the final experimental data are restricted to the tanks of two high-fat and three low-fat dietary groups with low mortality.

Wet mass (± 0.01 g) and fork length (± 1mm) of fish that were fasted for 24 h (August) or 48 hours (November) were measured, and fin clips of each individual were taken for future genetic identification analyses (see below), at two time points (in August and November). During the latter sampling, all fish were euthanized with an overdose of MS-222. Muscle tissue was selected for lipid analyses as it is known to be the site of an important fat depot in salmonid fishes (Jobling et al. 2002; Weil et al., 2013). A standardized muscle tissue sample from below the dorsal fin and above the lateral line was sampled and weighed for lipid extraction for 49 fish. These 49 samples were flash-frozen in liquid nitrogen and stored in −80 °C until analysis. In addition, the mass of total visceral fat tissue from the body cavity was measured (± 0.001 g) by scraping all visible fat carefully with a scalpel. Body condition was calculated as the deviation from the slope of logarithmic mass on logarithmic length - a correlate of salmon parr lipid content (Sutton et al., 2000). Visceral fat index (VFI) was calculated as total visceral fat mass divided by total fish mass multiplied by 100. All individuals were observed to be immature. This experiment was conducted under animal experimentation license -ESAVI/7248/04.10.07/2017.

### SNP genotyping

DNA was extracted from the fin clip samples using standard chelex or salt extraction methods and genotyped for 141 SNPs and a sexing marker as in Aykanat et al., (2016), except for modifications enabling genotyping on Illumina platform sequencers. These data were used to identity-match individuals from the two sampling events, in order to enable calculation of individual-specific growth rates, and to assign them to putative hatchery parents, to account for non-independence of data and for heritability calculations (see below), using the likelihood method implemented in SNPPIT 35 (Anderson, 2010). Parental genotypes were obtained from broodstock data generated in Debes et al., (2020b).

### Lipid extraction and chromatography

Muscle tissue was homogenized in Milli-Q water and the total lipids were extracted into chloroform according to Folch et al. (1957), and the extract solvent was evaporated with nitrogen stream. Immediately, the lipids were solved in 1.5 ml chloroform/methanol (1:2 vol/vol) and the sample solution was stored at −80° C until analysis. High Performance Thin Layer Chromatography was then used to separate and quantify concentrations of the main classes of polar lipids: phosphatidylcholine (PC), phosphatidylethanolamine (PE), phosphatidylserine (PS) and phosphatidylinositol (PI) and sphingomyelin (SM) eluted with a chloroform/methanol/acetic acid/water (25:17.5:3.8:1.75) solvent, and neutral lipids: cholesterol (C), triacylglycerol (TAG), and cholesterol ester (CE)eluted with a hexane/diethyl ether/acetic acid/water (26:6:0.4:0.1) solvent according to Lehti et al. (2018). Lipid concentrations were determined by scanning the charred lipid bands, created with acid and heating plate, of the samples on the silica plates under UV light at 254 nm (CAMAG TLC Scanner 3) and integrating peak areas using the winCATS program (CAMAG version 1.1.3.0) and comparing these to the lipid standards with known concentrations ranging from 50 pmol/μg to 60 pmol/μg. PS and PI were calculated under one peak as their peaks are very close to each other so individual concentrations for these two could not be calculated.

### Statistical Analysis

We fitted a series of multivariate linear animal models for focal traits (PC, PE, C, PS.PI, SM, CE, TAG, Condition, VFI) as responses to simultaneously test for fixed effects and estimate phenotypic correlations among response traits, adjusted for fixed and random effects, under REML using ASReml-R v. 3 (Butler et al., 2009) in R v. 3.0.2. Before analysis, we mean-centered and variance-standardized all traits except for Condition and VFI. As fixed effects we included terms for the diet (low- or high-fat diet), *vgll3* genotype (LL, LE, EE, where *L* and *E* indicate the alleles associated with <u>L</u>ate vs <u>E</u>arly maturation, respectively (Barson et al., 2015), and sex (male, female) and all possible interactions. As random terms we fitted tank effects (n = 5) and animal effects (n = 88 entries in the inverse of the additive relatedness matrix). Animal effects were included to account for the complex relatedness among the hatchery crossed individuals and thus possible genetic correlation for each trait. Such animal models ((Henderson, 1973, 1953) are also able to estimate the additive genetic variance accounted for by animal effects, but this may require a larger sample size for accurate estimates. To enable fitting the animal model, we constructed a pedigree from which we calculated the inverse of the additive genetic relatedness matrix. To test for fixed effects, we used *F*-test with denominator degrees of freedom approximated according to Kenward and Roger (1997).

## Results

### Heritability, and major locus effects, of lipid class concentrations, body condition, and VFI

Only two fish with the *vgll3*\*LL (late maturing) genotype were sampled, likely because the 49 individuals destined for lipid analysis were sampled randomly from the 673 individuals and the *vgll3*\*L allele is rare in the broodstock (Craig Primmer, unpublished data), which restricted the assessment of *vgll3* effects to a comparison of *vgll3*\**EE* and *vgll3*\**EL*. As *vgll3* effects were not detected in initial models, *vgll3* effects were removed from the final model. Also, as was expected with the available sample size, we did not detect significant additive genetic variance for lipid class concentrations. Nonetheless, additive genetic effects accounted for up to ~20% of the phenotypic variation (i.e., heritability, h^2^) for PS and PI (h^2^ values for each component in supplement). Heritability estimates ranged from 0 (C, CE, PE) to 0.2 (Condition), but twice the standard error of the estimate of all variables included zero, indicating that larger sample sizes would be needed in future studies (Supplementary Table A).

### Effects of feed treatment, sex, and body length

Interactions among fixed effects were not detected (not shown) and were, therefore, removed from the model. Furthermore, there were no detectable effects of diet, but sex and body length effects were detected (**Table 1**, **Figure 1**). The concentrations of the main storage lipid, TAG, the main structural lipid, PC, and the smaller lipid classes, C and SM, differed between males and females. Most traits (except for condition and VFI, which were already size standardized) covaried negatively with (Ln of) body length (**Table 1**, **Figure 1**).

**Table 1:** ANOVA table based on *F*-tests for the multivariate model including the non-significant feed treatment with lipid class concentrations being standardized for body mass (mol / g of tissue).

| Trait | Effect | Df | DDF | *F* | *P* |
| --- | --- | --- | --- | --- | --- |
| PC | Sex | 1 | 43.8 | 8.03 | **0.007** |
| PE | Sex | 1 | 44.0 | 2.49 | 0.122 |
| C | Sex | 1 | 44.0 | 5.50 | **0.024** |
| PS.PI | Sex | 1 | 44.3 | 0.77 | 0.384 |
| SM | Sex | 1 | 44.7 | 6.67 | **0.013** |
| TAG | Sex | 1 | 44.4 | 5.32 | **0.026** |
| CE | Sex | 1 | 43.5 | 0.59 | 0.448 |
| Condition | Sex | 1 | 43.9 | 2.65 | 0.111 |
| VFI | Sex | 1 | 40.7 | 0.08 | 0.782 |
| PC | Feed | 1 | 15.9 | 2.78 | 0.115 |
| PE | Feed | 1 | 44.0 | 0.59 | 0.446 |
| C | Feed | 1 | 44.0 | 1.06 | 0.309 |
| PS.PI | Feed | 1 | 9.5 | 2.24 | 0.167 |
| SM | Feed | 1 | 43.8 | 2.56 | 0.117 |
| TAG | Feed | 1 | 43.8 | 0.30 | 0.584 |
| CE | Feed | 1 | 11.6 | 0.00 | 0.973 |
| Condition | Feed | 1 | 2.4 | 0.01 | 0.920 |
| VFI | Feed | 1 | 2.1 | 4.42 | 0.166 |
| PC | ln.length | 1 | 43.7 | 21.82 | **< 0.001** |
| PE | ln.length | 1 | 44.0 | 4.58 | **0.038** |
| C | ln.length | 1 | 44.0 | 27.85 | **< 0.001** |
| PS.PI | ln.length | 1 | 41.9 | 6.59 | **0.014** |
| SM | ln.length | 1 | 43.2 | 12.10 | **0.001** |
| TAG | ln.length | 1 | 43.4 | 4.96 | **0.031** |
| CE | ln.length | 1 | 43.6 | 6.85 | **0.012** |
| Condition | ln.length | 1 | 35.7 | 0.00 | 0.981 |
| VFI | ln.length | 1 | 39.4 | 0.10 | 0.758 |

**Figure 1.**
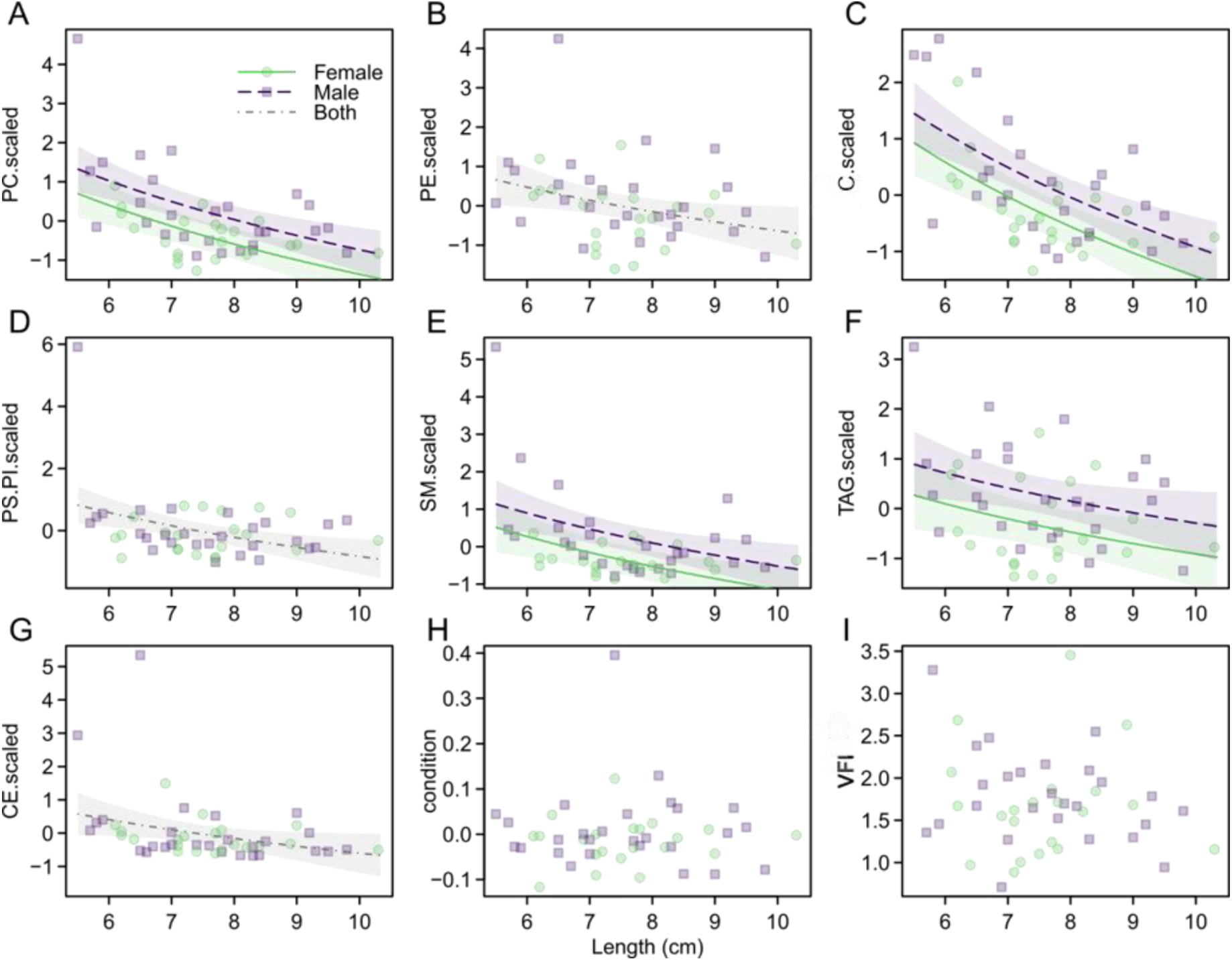
Sex effects and covariance with length for the focal traits as labelled in the Y-axis (individual scaled lipid class concentration; mol / g of tissue) vs. individual length (A-I). A detected covariance between a trait and length is presented as model-predicted regression line with 95% confidence bands. When sex effects were detected, the regression line is presented with separate elevations for males and females, commonly for both sexes otherwise.

### Correlations among lipid classes in muscle, body condition, and VFI

All lipid classes detected positively correlated with each other in our analyses (Figure 2). The main structural lipid class PC comprised ~23%, and the remaining membrane glycerophospholipids (, PE, PS, PI and SM) made up 26% of the total lipids detected. These four membrane glycerophospholipid classes positively correlated with PC, two of them significantly so, with correlation coefficients between 0.36 and 0.83. C significantly covaried with PC, PE, CE and SM, with correlation coefficients between 0.44 and 0.57. TAG also significantly covaried with all membrane structural lipid classes except C, with correlation coefficients between 0.45 and 0.76. PC, being the main component of lipid droplet surrounding monolayers, exhibited the strongest correlation with TAG. In addition, CE significantly covaried with three membrane structural lipid classes including PE, SM and C with correlation coefficients between 0.44 and 0.63. However, condition factor and VFI did not significantly co-vary with any lipid class concentration in muscle and many estimates, although not different from zero, were negative.

**Figure 2.**
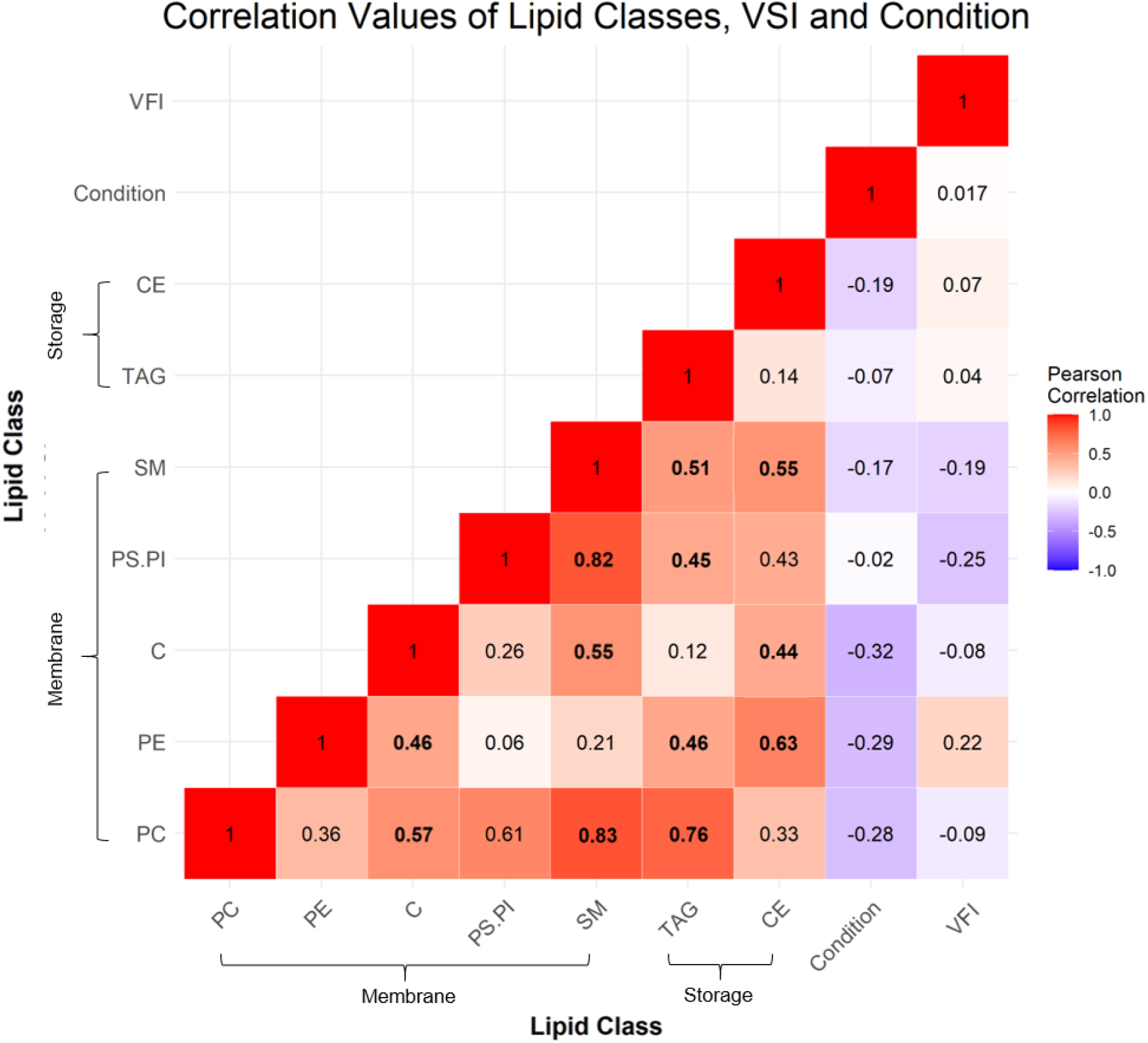
Pearson correlations between concentrations of lipid classes, condition factor, and VFI. All estimates are controlled for effects of lipid treatment, sex, body length, and experimental tank (i.e., partial correlations). Lipid classes are ordered from highest to lowest concentration within membrane lipids and highest to lowest concentrations within storage lipids. Red shows a positive correlation, whereas blue shows a negative correlation. Values in bold were significantly correlated (Bonferroni adjusted; Supplementary Table B).

## Discussion

We report here, for the first time to the best of our knowledge, that the muscle tissue of juvenile Atlantic salmon males had on average a higher concentration of several major lipid classes compared to females. We also observed a general decrease in lipid concentrations with increasing body length. Storage lipid concentrations, composed predominantly of TAG but also CE, also decreased with fish size. Lipid content is known to change based on many factors including fish size, feed ration and dietary fat level where major processes such as maturation or smoltification timing come into play (Hillestad and As, 1998; Nordgarden et al., 2003; Rowe et al., 1991; Sheridan, 1989; Storebakken and Austreng, 1987). These major factors could help explain the observed sex-specific differences in lipid class concentrations in the individuals. It is well known that there are sex-specific life-history differences as males and females can mature at different sizes and ages (Fleming, 1996). For example, faster growth increases the potential of maturation before sea migration, which is much more common in males than in females (Fleming, 1996). Starting reproduction during the juvenile phase is energetically costly (Myers, 1984; Whalen and Parrish, 1999). Therefore, juvenile males may require depositing larger stores of slowly mobilized energy in muscle than juvenile females. It has been found in many salmonid species, including Atlantic salmon, that during periods of increased growth there was also an increase in the β-oxidation capacity in muscle tissue resulting in lower lipid levels (Nordgarden et al., 2003). However, once males start maturing, growth decreases by two-fold compared to immature parr, which can lead to a delay of the size-dependent smoltification and thereby of the associated marine migration (Thorpe, 1986; Whalen & Parrish, 1999).

The timing of the marine migration is another life-history trait where levels of lipid reserves can potentially have an influence. It has been suggested that salmon juveniles can be separated into two groups based on growth (upper modal and lower modal) during the freshwater phase and the upper modal group will maintain a higher appetite and growth during winter months than the lower modal group (Higgins, 1985; Metcalfe, 1998; Morgan et al., 2002; Skilbrei, 1988; Thorpe, 1977) and therefore possibly maintain higher total body lipid reserves during winter (Higgins and Talbot, 1985). The upper modal group are more likely to smolt and migrate to the sea the following spring while the lower modal group are more likely to remain in freshwater for at least an additional year (Thorpe 1977, Bailey et al., 1980, Kristinsson et al., 1985). An earlier study showed that body length during the late summer and fall was able to predict the probability of a spring migration the following year and that both traits are genetically highly correlated (Debes et al., 2020a;). However, some individuals in the lower modal group can still grow rapidly during the winter and become migrants in the following spring (Debes et al., 2020a; Zydlewski et al., 2014).

Lipid-storing adipocytes are mainly located in the muscle and viscera; in the skeletal muscle, adipocytes largely reside in the myospeta, i.e., in the connective tissue between muscle fibers (Zhou et al., 1996). Visceral fat is assumed to provide much of the energy needed for the initiation of sexual maturation (Rowe et al., 1991). Fish on the high-fat diet had slightly more stored visceral fat as measured by VFI than those on the low-fat diet (however, this result is not very strongly supported, see above, Table 1). This is expected as it has been found that fish store more visceral fat with increasing lipid and carbohydrate levels in feed, whereas muscle fat can be reduced with increasing protein levels in feed (Aksnes, 1995; Hillestad and As, 1998; Weihe et al., 2019). However, no significant correlation between VFI and any muscle lipid class was detected, nor were there significant VFI differences detected between the diet treatments. The relatively small difference in visceral fat levels between the high and low-fat diet treatments may suggest that the difference in lipid content between the feeds (12.6% vs 19.9%) was not sufficiently large to give a significant difference for the current sample sizes. Faster growing fish may not store more lipid, relative to their total mass, but start to store lipids in different tissues throughout the body. For example, males may potentially deposit lipids in the muscles, while female may deposit more lipids in the viscera, but further research is required to test this hypothesis. Behavioral experiments with Masu salmon *Oncorhynchus masou* fry provide one explanation for the male’s larger investment of lipid to muscle, since more males were able to swim upstream, which may be due to genetically determined differences in muscle energetics linked with sex (Nagata and Irvine, 1997).

Investigating muscle lipid class profiles, and identifying sex-specific patterns, as we did here, contributes to understanding of energy allocation, differential physiological performance and behavior among individuals and, ultimately, survival of juvenile Atlantic salmon (Næsje et al., 2006; Post & Parkinson, 2001). Here, we presented evidence in Atlantic salmon that i) males have higher concentrations of membrane and storage lipids in skeletal muscle than females and ii) the muscle lipid concentrations covaried negatively with body length. Future studies would benefit from greater samples sizes, more specific lipid species identification, including additional tissue types, to better understand the importance of controlling body lipid distribution in juvenile fish preparing for challenging future life history events.

## Acknowledgements

We thank Raisioaqua, and in particular Susanna Airaksinen and Thomas Ginström for arranging the low-fat feed for the experiment. We further thank Nico Lorenzen, Suvi Ikonen and Thomas van der Geest for their help in husbandry and sampling of the fish, Jacqueline Moustakas-Verho for discussions and manuscript comments, Annukka Ruokolainen, Noora Parre, and Seija Tillanen for their help in the genetics lab and Hanna Ruhanen, Sanna Sihvo and Minna Holopainen for their help in the lipidomics lab and for discussions.

## Appendix

**Table A.** Summary of h^2^ estimates with standard errors for all traits for the 49 individuals.

| Trait | Estimate | SE |
| --- | --- | --- |
| VFI | 0.06 | 0.26 |
| Condition | 0.21 | 0.36 |
| TAG | 0.05 | 0.08 |
| C | 0.00 | 0.00 |
| CE | 0.00 | 0.00 |
| SM | 0.08 | 0.08 |
| PE | 0.00 | 0.00 |
| PS.PI | 0.09 | 0.17 |
| PC | 0.01 | 0.06 |

**Table B.**
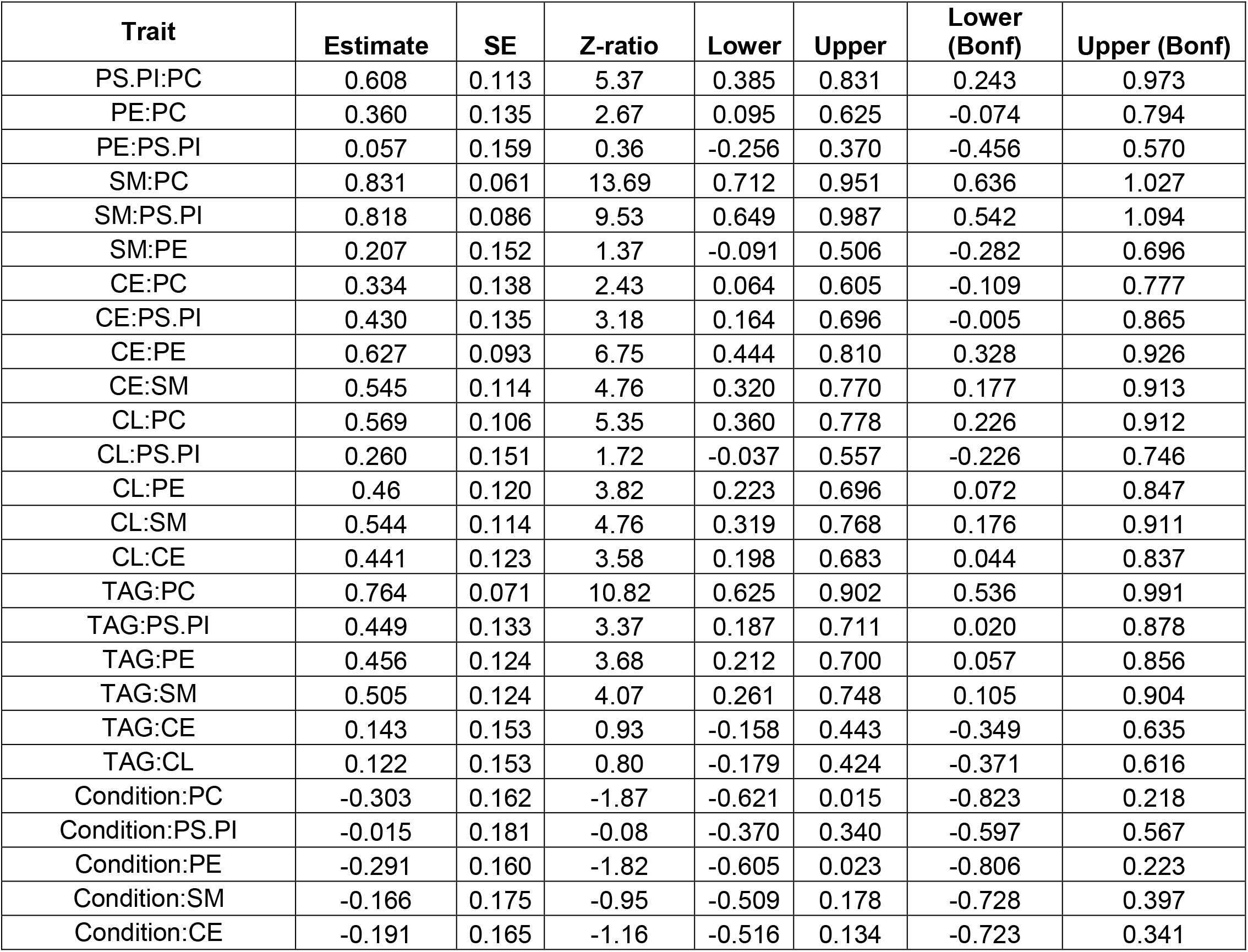
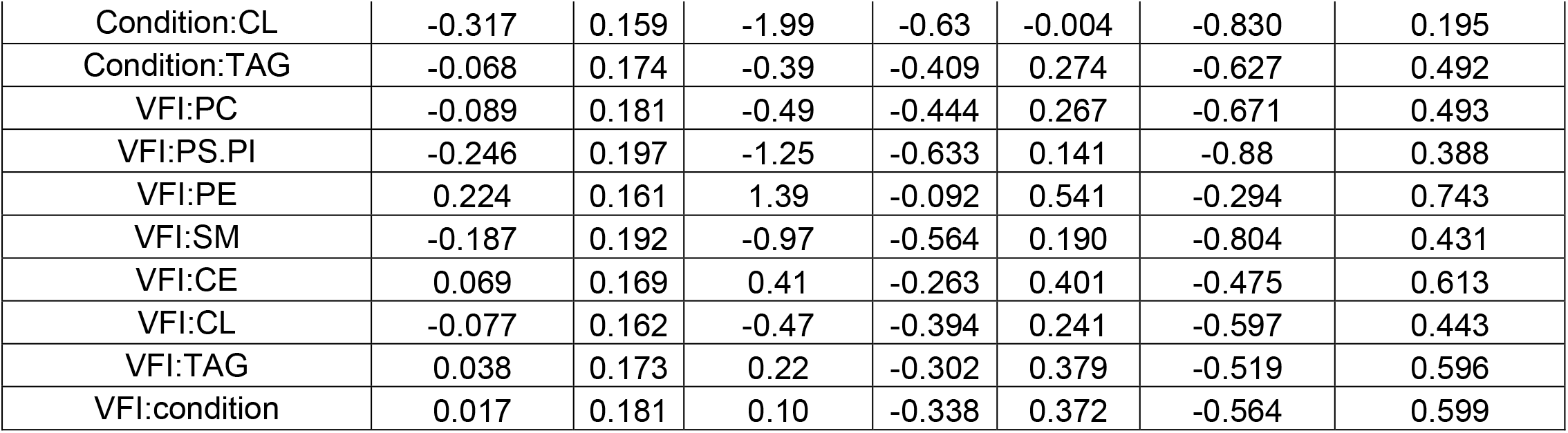
Summary of correlation estimates with standard errors and raw and Bonferroni-corrected lower and upper 95% confidence intervals.

